# Massively differential bias between two widely used Illumina library preparation methods for small RNA sequencing

**DOI:** 10.1101/001479

**Authors:** Jeanette Baran-Gale, Michael R. Erdos, Christina Sison, Alice Young, Emily E. Fannin, Peter S. Chines, Praveen Sethupathy

## Abstract

Recent advances in sequencing technology have helped unveil the unexpected complexity and diversity of small RNAs. A critical step in small RNA library preparation for sequencing is the ligation of adapter sequences to both the 5’ and 3’ ends of small RNAs. Two widely used protocols for small RNA library preparation, Illumina v1.5 and Illumina TruSeq, use different pairs of adapter sequences. In this study, we compare the results of small RNA-sequencing between v1.5 and TruSeq and observe a striking differential bias. Nearly 100 highly expressed microRNAs (miRNAs) are >5-fold differentially detected and 48 miRNAs are >10-fold differentially detected between the two methods of library preparation. In fact, some miRNAs, such as miR-24-3p, are over 30-fold differentially detected. The results are reproducible across different sequencing centers (NIH and UNC) and both major Illumina sequencing platforms, GAIIx and HiSeq. While some level of bias in library preparation is not surprising, the apparent massive differential bias between these two widely used adapter sets is not well appreciated. As increasingly more laboratories transition to the newer TruSeq-based library preparation for small RNAs, researchers should be aware of the extent to which the results may differ from previously published results using v1.5.

## Introduction

Small RNAs, such as microRNAs (miRNAs), are important regulators of gene expression in a wide variety of normal biological and pathological processes(1,2). Numerous technologies, including quantitative PCR (qPCR), microarray, and deep sequencing, are presently in use for high-throughput miRNA profiling(3–5). Though each of these methods has both advantages and limitations, deep sequencing has emerged as the gold standard for discovery and quantification of miRNAs, particularly for those that are of low abundance. There are several small RNA sequencing technologies, including Illumina, Applied Biosystems (ABI) SOLiD, and 454 Life Sciences that are commercially available. Recent studies have demonstrated that each of these technologies harbors different limitations that lead to variable biases(6,7).

A critical step in the preparation of a small RNA library for deep sequencing is the ligation of adapter sequences to both ends of small RNAs. These adapters provide the template for primer-based reverse transcription, amplification, and sequencing. The efficiency of adapter ligation is thought to depend on the adapter sequence, the ligase, as well as the nucleotide composition and secondary structures of the small RNAs to be sequenced(8,9). Illumina introduced the v1.5 small RNA library preparation method in February 2009 and the TruSeq method more recently in November 2010. Because one critical difference between these methods is the adapter sequences, some level of differential bias between these two methods is expected. However, the extent of the bias has not been evaluated previously and could be important for guiding accurate comparison of miRNA expression results between these two methods.

In this study, we directly compared the small RNA sequencing results between Illumina v1.5 and TruSeq. We also performed the sequencing on two different Illumina platforms (GAIIx and HiSeq) and at two different sequencing centers (UNC and NIH). While we expected some level of bias in the library preparation, the apparent extensive differential bias between these two widely used Illumina adapter sets is striking and not reported previously. For example, nearly 25 of the most abundant miRNAs are >10-fold differentially detected between v1.5 and TruSeq. This finding serves as an important caution, particularly to laboratories/facilities that used v1.5 but are now transitioning to the newer TruSeq protocol. In general, the findings of this study add to the growing body of literature on bias in small RNA sequencing that merits continued investigation, particularly with regard to the development of strategies for bias remediation.

## Materials and Methods

### Sequencing and bioinformatic analysis

Mouse insulinoma (MIN6) cells were cultured as previously described(10). Cells were lysed and RNA was isolated using either the Norgen Total RNA Purification Kit (UNC) or TRIzol-mediated extraction (NIH). Only samples with an RNA Integrity Number (RIN) of 8.5 or higher, as measured by Agilent Bioanalyzer 2100, were considered for further analysis. Small RNA libraries were generated using either the Illumina v1.5 protocol or the Illumina TruSeq protocol. Sequencing was performed on either the Illumina GAIIx or Illumina HiSeq 2000 platforms. Small RNA-seq reads were trimmed using cutAdapt (-O 10 –e 0.1) to remove remnants of the 3’-adapter sequence. Subsequent mapping of trimmed reads to the mouse genome and miRNA/isomiR quantification were performed exactly as previously described(10).

### Real time quantitative PCR analysis

MIN6 cells were cultured and lysed as above and RNA was isolated using the Norgen Total RNA Purification Kit. Complementary DNA (cDNA) was synthesized using the TaqMan miRNA Reverse Transcription kit (Applied Biosystems; Grand Island, NY) according to the manufacturer’s instructions. Real-time PCR amplification was performed using TaqMan miRNA assays in TaqMan Universal PCR Master Mix on a BioRad CFX96 Touch Real Time PCR Detection system (Bio-Rad Laboratories, Inc., Richmond, CA). Reactions were performed in triplicate using *U6* as the internal control. miRNA levels were expressed as relative quantitative values, which represent fold differences relative to miR-30e-5p. All TaqMan assays used in this study where purchased from Applied Biosystems, Inc. (Grand Island, NY) and include: mmu-miR-24-3p (4427975-000402), mmu-miR-27b-3p (4427975-000409), mmu-miR-29a-3p (4427975-002112), mmu-miR-375-3p (4427975-000564), miR-30e-5p (4427975-002223), and *U6* (4427975-001973).

## Results

We isolated RNA from a widely-used pancreatic beta-cell-like cell line (MIN6) and performed small RNA-seq according to three different methods: [1] Illumina v1.5 library preparation sequenced on GAIIx platform (v1.5-GAIIx), [2] Illumina TruSeq library preparation sequenced on GAIIx platform (TS-GAIIx), and [3] Illumina TruSeq library preparation sequenced on HiSeq platform (TS-HiSeq). TS-GAIIx and v1.5GAIIx were carried out at the NIH Intramural Sequencing Center (NISC) on June 25^th^, 2013 and TS-HiSeq was performed at the UNC High throughout Sequencing Facility (HTSF) on June 6^th^, 2013. Three replicate small RNA libraries were generated for each method, yielding a total of nine small RNA-seq datasets.

We used our previously published bioinformatic pipeline(10) to analyze the small RNA-seq reads in each dataset. Results of the 3’-adapter trimming and genome mapping are provided in Table S1. Total number of reads across the nine datasets range from ∼17 million to ∼24 million (Table S1). In each of the datasets, >70% of the align-able reads map to annotated miRNAs and >1000 distinct mature miRNAs are represented by at least ten reads. Among these miRNAs, 315 have a relative expression of at least 100 reads per million mapped reads (RPMM) in at least one library (Table S2). We refer to these miRNAs as “highly expressed.” To compare miRNA expression profiles across datasets, we correlated the expression profiles of these abundant miRNAs across all nine datasets.

The miRNA expression profiles from replicates within each method are very highly correlated (r^2^ > 0.99), clearly demonstrating that both the method of library preparation and the sequencing platform yield exceptionally reproducible results (Figure 1). Furthermore, we also observed a very strong correlation (r^2^ > 0.90) among TS-GAIIx and TS-HiSeq samples, but substantially lower correlation (r^2^ ∼ 0.6) among TS-GAIIx (or TS-HiSeq) and v1.5-GAIIx samples (Figure 1). These results indicate that neither sequencing platform (GAIIx vs. HiSeq) nor sequencing facility (UNC vs. NIH) is a major contributor to technical variation, but that the method of library preparation (TS vs. v1.5) is a significant factor.

**Figure 1.**
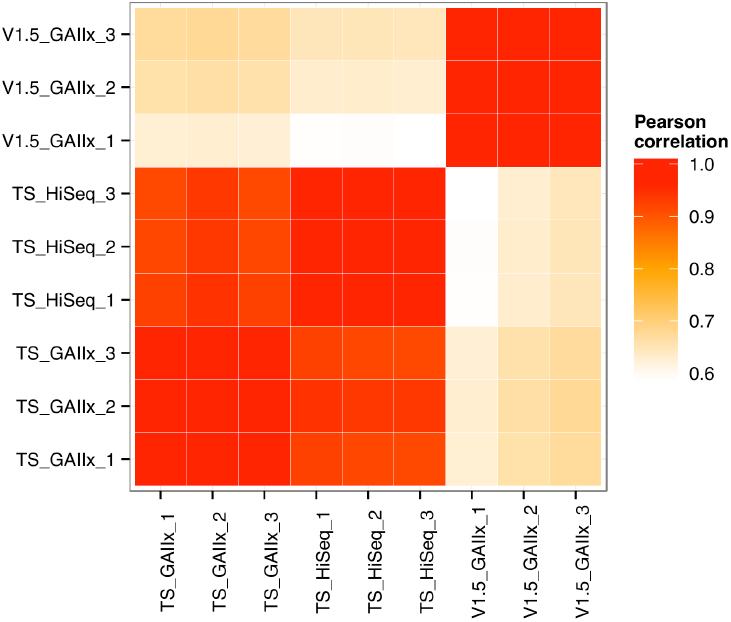
Comparison of miRNA expression profiles between two different Illumina library preparation protocols reveals massive differential bias. A comparison of the following three methods is shown: Illumina v1.5 library preparation sequenced on GAIIx platform (v1.5_GAIIx), Illumina TruSeq library preparation sequenced on GAIIx platform (TS_GAIIx), and Illumina TruSeq library preparation sequenced on HiSeq platform (TS_HiSeq). Three replicate small RNA libraries were generated for each of the three methods. Correlation values were calculated by Pearson’s metric. Similar results were obtained based on the calculation of Spearman’s correlation coefficient (rho). Red and white colors indicate strongest and weakest correlation, respectively.

Among the 315 highly expressed miRNAs included in the correlation analysis, only 8 are >10-fold differentially detected between TS-GAIIx and TS-HiSeq (Figure 2A). Moreover, most of these differentially detected miRNAs are on the lower end of the expression spectrum (Table S2). In stark contrast, when comparing TS-GAIIx with v1.5-GAIIx, 48 are >10-fold differentially detected and 96 are >5-fold differentially detected (Figure 2B). Strikingly, ∼79% (n=38/48) of the former and ∼70% (n=67/96) of the latter set of miRNAs are present at greater abundance in the samples prepared by v1.5 compared to the samples prepared by TruSeq (Figure 2B). These miRNAs include several that are known regulators of beta cell development and function, including miR-24-3p(11), miR-29b-3p(12), and miR-200c-3p(13), which are ∼36-fold, ∼31-fold, and ∼13-fold more highly detected in the samples prepared by v1.5, respectively. For example, miR-24-3p is among the top ten most highly expressed miRNAs in MIN6 cells according to v1.5, but is consistently not even in the top hundred according to TruSeq (Table S2). It is worth noting that despite the overall bias toward higher miRNA expression levels in samples prepared by v1.5, a few miRNAs are more highly detected in samples prepared by TruSeq (Figure 2B). For example, miR-26a-5p, which is known to have functional relevance in the beta cell(14), is among the top ten most highly expressed miRNAs in MIN6 cells according to TruSeq, but is scarcely in the top fifty according to v1.5 (Table S2).

**Figure 2.**
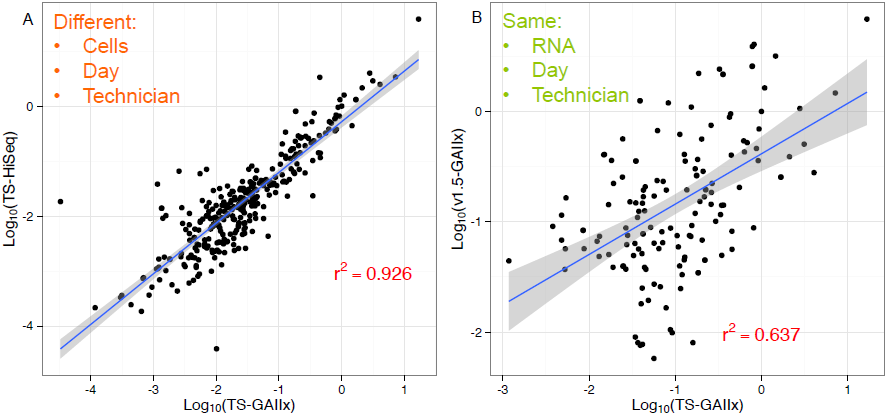
Nearly fifty of the most abundant miRNAs are greater than ten-fold differentially detected between Illumina v1.5 and TruSeq. (A) Relative expression levels of the most abundant (n=315) miRNAs according to the GAIIx and HiSeq sequencing platforms are shown. **(B)** Relative expression levels of the most abundant (n=315) miRNAs according to the v1.5 and TruSeq (TS) library preparation methods are shown. Relative miRNA expression levels were calculated according to the following: log_10_(miRNA RPMM / miR-30e-5p RPMM), where RPMM is reads per million mapped reads. miR-30e-5p represents a housekeeping miRNA, due to its invariance and robust expression across all datasets. Similar results were obtained from using another invariant but lowly expressed housekeeping miRNA, miR-130b.

**Figure 3.**
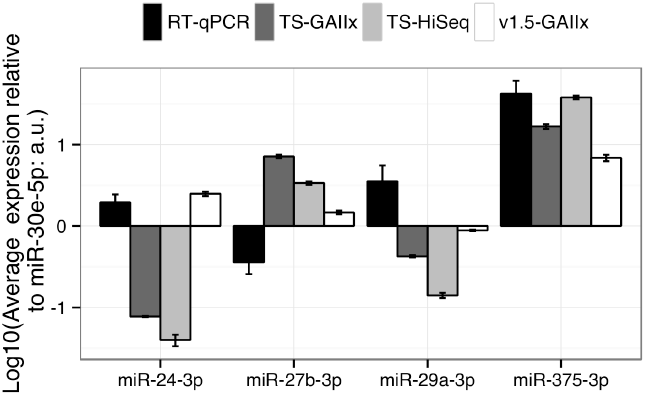
Results from quantitative PCR are not uniformly consistent with one method of library preparation. Comparison of relative expression levels for four miRNAs (miR-24-3p, miR-27b-3p, miR-29a-3p, and miR-375-3p) across four different methods of miRNA detection are shown. miRNA expression levels were normalized to miR-30e-5p, which represents a housekeeping miRNA due to its invariance and robust expression across all datasets.

We selected four of the highly expressed miRNAs, miR-24-3p, miR-27b-3p, miR-29a-3p, and miR-375-3p for quantification by TaqMan-based real time quantitative PCR (RT-qPCR). To facilitate a comparison of the findings between RT-qPCR and the sequencing methods, we normalized the expression levels of each miRNA to that of miR-30e-5p, which is highly expressed and invariant across all of the datasets. The qPCR result matches that of v1.5 for miR-24-3p, but more closely resembles that of TruSeq for miR-375 and is completely different than both v1.5 and TruSeq for miR-27b-3p and miR-29a-3p. These results indicate that qPCR is not uniformly consistent with one method of library preparation for sequencing.

## Discussion

The presence of bias in small RNA profiling is well established in the literature(8,15–17). Differential bias across various expression platforms (e.g., microarray, qPCR, sequencing) and sequencing technologies (e.g., Illumina, ABI SOLiD, 454 Life Sciences) has also been demonstrated(6,7,18). However, no study has focused on different library preparation methods within the same sequencing technology. Here we compare two of the most popular methods from Illumina (v1.5 and TruSeq). The results of our study point to a massive differential miRNA detection bias between these two library preparation methods. This finding was independent of the sequencing center (NIH, UNC) and sequencing platform (GAIIx, HiSeq). While some level of bias in library preparation is not surprising, the apparent extensive differential bias between these two widely used adapter sets is striking and not well appreciated.

It is important to note that our study does not suggest that one method of library preparation is necessarily always more reliable or accurate for miRNA detection than the other; rather, it likely depends on the miRNA. Specifically, the ligation efficiencies of different adapter sequences may differ based on features that vary across miRNAs, such as nucleotide sequence, chemical modification, and secondary structure(8,9,19). The factors that control the differential biases specifically between the adapter sets used in v1.5 and TruSeq merit further detailed investigation. Researchers seeking to ameliorate the influence of these biases on miRNA expression levels can consider using pools of different adapter sets during library preparation(20,21) or alternative RNA cloning methods(22).

As increasingly more laboratories transition to the newer TruSeq-based library preparation for small RNAs, researchers should be aware of the extent to which the results may differ from previously published results using the v1.5 method. We strongly caution researchers against merging together small RNA-seq data generated from v1.5 and TruSeq library preps. Also, in any standard small RNA-seq study, in which only one adapter set is used for library preparation, one should be aware of the potential pitfalls of applying arbitrary cutoffs based on expression (such as “top 100 detected”) to identify miRNAs for further functional analysis, because some miRNAs that appear lowly expressed could be inefficiently detected for purely technical reasons (such as miR-24 in the TruSeq datasets presented in this study). In general, we recommend against using small RNA-seq data to make calls on the “absolute” levels of miRNAs, unless additional precaution has been taken to substantially mitigate the biases discussed here. Despite these issues, deep sequencing is still an extremely valuable method for *de novo* discovery of isomiRs and novel small RNAs, as well as for studying relative miRNA expression changes across different conditions or time points.

## Data Availability

MIN6 TS_HiSeq reads are available at Gene Expression Omnibus (GEO) under the accession number GSE44262. MIN6 v1.5_GAIIx and TS_GAIIx reads are will also be made available at GEO (submission in progress).

## Supplementary Material

Supplementary Data are available online at http://biorxiv.org/.

## Acknowledgments

The authors wish to thank members of the Sethupathy laboratory for critical discussions and technical suggestions.

## References

1. Bartel, D.P. (2009) MicroRNAs: target recognition and regulatory functions. Cell, 136, 215–233.

2. Couzin, J. (2008) MicroRNAs make big impression in disease after disease. Science, 319, 1782–1784.

3. Gunaratne, P.H., Coarfa, C., Soibam, B. and Tandon, A. (2012) miRNA data analysis: next-gen sequencing. Methods Mol Biol, 822, 273–288.

4. Baker, M. (2010) MicroRNA profiling: separating signal from noise. Nature methods, 7, 687–692.

5. Pritchard, C.C., Cheng, H.H. and Tewari, M. (2012) MicroRNA profiling: approaches and considerations. Nature reviews. Genetics, 13, 358–369.

6. Tian, G., Yin, X., Luo, H., Xu, X., Bolund, L., Zhang, X., Gan, S.Q. and Li, N. (2010) Sequencing bias: comparison of different protocols of microRNA library construction. BMC biotechnology, 10, 64.

7. Linsen, S.E., de Wit, E., Janssens, G., Heater, S., Chapman, L., Parkin, R.K., Fritz, B., Wyman, S.K., de Bruijn, E., Voest, E.E. et al. (2009) Limitations and possibilities of small RNA digital gene expression profiling. Nature methods, 6, 474–476.

8. Hafner, M., Renwick, N., Brown, M., Mihailovic, A., Holoch, D., Lin, C., Pena, J.T., Nusbaum, J.D., Morozov, P., Ludwig, J. et al. (2011) RNA-ligase-dependent biases in miRNA representation in deep-sequenced small RNA cDNA libraries. RNA, 17, 1697–1712.

9. Zhuang, F., Fuchs, R.T., Sun, Z., Zheng, Y. and Robb, G.B. (2012) Structural bias in T4 RNA ligase-mediated 3’-adapter ligation. Nucleic acids research, 40, e54.

10. Baran-Gale, J., Fannin, E.E., Kurtz, C.L. and Sethupathy, P. (2013) Beta Cell 5’-Shifted isomiRs Are Candidate Regulatory Hubs in Type 2 Diabetes. PloS one, 8, e73240.

11. Zhu, Y., You, W., Wang, H., Li, Y., Qiao, N., Shi, Y., Zhang, C., Bleich, D. and Han, X. (2013) MicroRNA-24/MODY gene regulatory pathway mediates pancreatic beta-cell dysfunction. Diabetes, 62, 3194–3206.

12. Pullen, T.J., da Silva Xavier, G., Kelsey, G. and Rutter, G.A. (2011) miR-29a and miR-29b contribute to pancreatic beta-cell-specific silencing of monocarboxylate transporter 1 (Mct1). Molecular and cellular biology, 31, 3182–3194.

13. Liao, X., Xue, H., Wang, Y.C., Nazor, K.L., Guo, S., Trivedi, N., Peterson, S.E., Liu, Y., Loring, J.F. and Laurent, L.C. (2013) Matched miRNA and mRNA signatures from an hESC-based in vitro model of pancreatic differentiation reveal novel regulatory interactions. Journal of cell science, 126, 3848–3861.

14. Melkman-Zehavi, T., Oren, R., Kredo-Russo, S., Shapira, T., Mandelbaum, A.D., Rivkin, N., Nir, T., Lennox, K.A., Behlke, M.A., Dor Y., et al. (2011) miRNAs control insulin content in pancreatic beta-cells via downregulation of transcriptional repressors. The EMBO journal, 30, 835–845.

15. Alon, S., Vigneault, F., Eminaga, S., Christodoulou, D.C., Seidman, J.G., Church, G.M. and Eisenberg, E. (2011) Barcoding bias in high-throughput multiplex sequencing of miRNA. Genome research, 21, 1506–1511.

16. Van Nieuwerburgh, F., Soetaert, S., Podshivalova, K., Ay-Lin Wang, E., Schaffer, L., Deforce, D., Salomon, D.R., Head, S.R. and Ordoukhanian, P. (2011) Quantitative bias in Illumina TruSeq and a novel post amplification barcoding strategy for multiplexed DNA and small RNA deep sequencing. PloS one, 6, e26969.

17. Linsen, S.E. and Cuppen, E. (2012) Methods for small RNA preparation for digital gene expression profiling by next-generation sequencing. Methods Mol Biol, 822, 205–217.

18. Willenbrock, H., Salomon, J., Sokilde, R., Barken, K.B., Hansen, T.N., Nielsen, F.C., Moller, S. and Litman, T. (2009) Quantitative miRNA expression analysis: comparing microarrays with next-generation sequencing. RNA, 15, 2028–2034.

19. Sorefan, K., Pais, H., Hall, A.E., Kozomara, A., Griffiths-Jones, S., Moulton, V. and Dalmay, T. (2012) Reducing ligation bias of small RNAs in libraries for next generation sequencing. Silence, 3, 4.

20. Sun, G., Wu, X., Wang, J., Li, H., Li, X., Gao, H., Rossi, J. and Yen, Y. (2011) A bias-reducing strategy in profiling small RNAs using Solexa. RNA, 17, 2256–2262.

21. Jayaprakash, A.D., Jabado, O., Brown, B.D. and Sachidanandam, R. (2011) Identification and remediation of biases in the activity of RNA ligases in small-RNA deep sequencing. Nucleic acids research, 39, e141.

22. Zhang, Z., Lee, J.E., Riemondy, K., Anderson, E.M. and Yi, R. (2013) High-efficiency RNA cloning enables accurate quantification of miRNA expression by deep sequencing. Genome biology, 14, R109.

